# Temporal Propagation of Neural State Boundaries in Naturalistic Context

**DOI:** 10.1101/2025.02.28.640737

**Authors:** Djamari Oetringer, Sarah Henderson, Dora Gözükara, Linda Geerligs

## Abstract

Our senses receive a continuous stream of complex information, which we segment into discrete events. Previous research has related such events to neural states: temporally and regionally specific stable patterns of brain activity. The aim of this paper was to investigate whether there was evidence for top-down or bottom-up propagation of neural state boundaries. To do so, we used intracranial measurements with high temporal resolution while subjects were watching a movie. As this is the first study of neural states in intracranial data in the context of event segmentation, we also investigated whether known properties of neural states could be replicated. The neural state boundaries indeed aligned with stimulus features and between brain areas. Importantly, we found evidence for top-down propagation of neural state boundaries at the onsets and offsets of clauses. Interestingly, we did not observe a consistent top-down or bottom-up propagation in general across all timepoints, suggesting that neural state boundaries could propagate in both a top-down and bottomup manner, with the direction depending on the stimulus input at that moment. Taken together, our findings provide new insights on how neural state boundaries are shared across brain regions and strengthen the foundation of studying neural states in electrophysiology.

## Introduction

The continuous influx of sensory input in our daily lives encompasses a vast amount of information. Despite its complexity, we can effortlessly process and utilize this information. This is due, in part, to our ability to automatically segment information into meaningful units (Magliano et al., 2014; Zacks, 2020; Zacks et al., 2007), which supports information processing (Kurby and Zacks, 2008), as well as memory encoding and retrieval (Bird, 2020; Gold et al., 2017; Sargent et al., 2013; Shin and DuBrow, 2020; Swallow et al., 2009). Recently, various studies have investigated neural states as a possible neural underpinning of event segmentation. Neural states are temporarily stable patterns of brain activity in a local brain area, sometimes also termed “events in the brain” or “neural events”, which are observed across the cortex and are usually studied while subjects experience a naturalistic stimulus (Baldassano et al., 2017; Geerligs et al., 2022; Oetringer et al., 2025; Sava-Segal et al., 2023).

Although it has been shown that transitions in neural states (i.e., neural state boundaries) in different brain regions co-occur forming a nested hierarchy (Baldassano et al., 2017; Geerligs et al., 2022), it is not known how neural states are propagated across the cortex and what the role of bottom-up versus top-down signaling is. Looking at event segmentation, there is substantial evidence that both major low-level perceptual changes (Heusser et al., 2018) as well as changes in high-level features of a narrative, such as a change in the goals of a character (Cutting, 2014; Zacks et al., 2009), can result in the perception of an event boundary. Similarly, both low-level and high-level features can lead to co-occurring neural state boundaries in feature-relevant areas (Oetringer et al., 2025). Given our previous observation of the nested hierarchy of boundaries, this suggests that boundaries could propagate across the cortex in a bottom-up but also in a topdown manner. For example, when there is a deviation from a social script that is detected in a higher-level brain area, such as a person introducing themselves with a high five rather than a handshake, this may result in a change in expectation for upcoming perceptual input, resulting in a neural state boundary in a lower-level area. Alternatively, a strong change in perceptual features could induce a boundary in a low-level area, which could then propagate to a high-level area when that perceptual change also results in changes in high-level representations of the scene.

Of particular interest when studying different levels of information and their interaction in the context of event segmentation is naturalistic language, which forms a nested hierarchy akin to events and neural states: multiple phonemes together form words, words form sentences, and sentences form narratives. Lexical processing is more apparent in lower-level auditory areas (e.g., anterior superior temporal gyrus), whereas the more complex syntactic integrations are more evident in higher-level areas (e.g., inferior frontal gyrus; Friederici, 2002). Although less rich in sensory information, reading narratives also results in transient increases in activity around event boundaries in a network of brain regions that is similar to those found when watching audiovisual movies (Speer et al., 2007). Indeed, when reading a narrative, readers perceive temporal shifts as event boundaries (Speer and Zacks, 2005). Large language models such as GPT-2 and GPT-3 have also been found to be able to segment text-only narratives similar to human event segmentation by providing instructions as one would do with human subjects (Michelmann et al., 2025), or by extracting measures of surprise and comparing those to the timings of human-annotated event boundaries (Kumar et al., 2023).

One way to infer the presence of top-down and bottom-up propagation of neural state boundaries is by investigating timing differences between neural states in different brain regions and in relation to particular stimulus features. Studies so far have been unable to investigate these timings as they were mostly using fMRI, which has very high spatial resolution but low temporal resolution. Instead, an electrophysiological method would be necessary to study the exact timings of neural state boundaries. Some recent studies have already explored the notion of neural states in electrophysiological data. In particular, Mariola et al. (2022) showed that neural states can be extracted from electroencephalogram (EEG) data, and that neural state boundaries align with event boundaries in the stimulus. Silva et al. (2019) related neural states in EEG to memory, by showing that neural state patterns during movie watching were reactivated during free recall of that movie. Finally, Henderson et al. (2025) further investigated the relationship between neural states and memory in different age groups, and found that greater distinction between neural states relates to better subsequent memory performance independent of age. Taken together, these studies show that neural states in electrophysiological methods show at least some similar characteristics as found in fMRI and, more importantly, could offer valuable insights into neural states that may not be possible in fMRI.

In this study, we intended to study the timing differences between neural state boundaries and the stimulus, and between the neural states of multiple brain areas. To this end, we used electrocorticography (ECoG), because, unlike EEG and MEG, it avoids issues of low spatial specificity by measuring local signals directly from different brain regions (Dubey and Ray, 2019). As we were interested in natural language comprehension, we looked at neural states in a high-level and a low-level language area in an open intracranial electroencephalography (iEEG) dataset, which included ECoG data, and in which participants watched a movie. To first test whether neural states can be found and studied in ECoG data, we aimed to replicate neural state features observed in fMRI. We tested whether neural state boundaries in ECoG data also align with stimulus features, as well as align between different brain areas, and whether they show the expected temporal hierarchy of high-level areas exhibiting longer neural states than low-level areas. We hypothesized that these three strongly supported findings in fMRI are found in ECoG data as well, and that a topdown and/or bottom-up flow of information would be visible in the exact timing of the neural state boundaries. More specifically, Chang et al. (2022) reported a bottom-up between-network lag gradient in naturalistic language processing that was absent during rest, which is why we expected a bottom-up propagation of boundaries in general. However, although bottom-up processes could be dominant, others have reported the presence of top-down predictions during natural language processing in MEG (Heilbron et al., 2022). Thus, we also expected to see top-down propagation at moments in time that are important for high-level processing, such as the end of clauses. To preface our results, we indeed replicated the finding that neural states align with stimulus features and between brain areas. However, we did not find that neural states in higher-level areas are longer than those in lower-level areas, which was unexpected given the duration hierarchy observed in fMRI data. In addition, we found evidence for a top-down flow of information by looking at the exact timing of neural state boundaries in different areas in relation to the stimulus.

## Results

To study top-down and bottom-up propagation of neural state boundaries, we identified neural states in an ECoG dataset, in which participants watched a short audiovisual movie. Specifically, we focused on brain activity in a low-level language region of interest (ROI; Brodmann areas 20, 21, 22, 41, 42) and a high-level language ROI (Brodmann areas 38, 39, 40, 44, 45, 46, 47). Per subject, each ROI contained at least 12 electrodes. One subject however had no electrodes in the low-level ROI and was therefore excluded from any analyses that required this ROI. See Supplementary Section A for the number of electrodes per subject per ROI. To extract neural state boundary timings from these ROIs, we applied Greedy State Boundary Search (GSBS; Geerligs et al., 2021) to single-subject data per ROI. Using GSBS, we computed the optimal number of states per subject per ROI in a fully data-driven manner by using an adjusted t-distance metric. We used annotations of word and clause on- and offsets to link the neural state boundaries to stimulus features. Because this is the first study of neural state boundaries in ECoG data, we first set out to replicate the findings of previous fMRI studies. Specifically, we investigated whether boundaries align with stimulus features, whether they overlap across brain regions, and whether a duration hierarchy is present. Next, we investigated whether there was any evidence of top-down and bottom-up propagation of neural state boundaries. To do so, we investigated the timing differences across regions in their alignment to the stimulus and to each other.

### Neural state boundaries align with speech

To assess whether neural states in ECoG data are related to the stimulus, we investigated the alignment between neural state boundaries and stimulus features, specifically the on- and offsets of words and clauses. In the low-level ROI, subjects had an average of 236.7 neural state boundaries across 3 minutes (standard deviation of 130.5) and 285.1 boundaries in the high-level ROI (standard deviation of 140.1), which are both much lower than the number of onsets and offsets of words (538), and higher than the number of onsets and offsets of clauses (152). See Supplementary Section B for example visualizations of the timelines. Per ROI and per stimulus feature, we computed the maximum Gaussian match relative to chance-level between the neural state boundaries in the ROI and the onsets and offsets of the stimulus feature. The Gaussian match is a non-commutative measure for how well boundaries in one timeline (e.g., the onsets and offsets of words) align to boundaries in another timeline (e.g., neural state boundaries in a specific ROI). Because of the high temporal resolution of ECoG, the Gaussian match was first computed across multiple possible delays, from which the maximum was taken to get the maximum Gaussian match. Using a one-tailed Wilcoxon signed-rank test on the maximum relative Gaussian matches per subject (N = 11 for high-level ROI, N = 10 for low-level ROI), we indeed found a significant match between neural states and the on- and offsets of clauses (Figure 1A; low-level ROI: *p* < 0.001, mean = 0.072, SD = 0.049; high-level ROI: *p* = 0.034, mean = 0.036, SD = 0.045). For words, both the low-level ROI (*p* = 0.065, mean = 0.005, SD = 0.049) and the high-level ROI (*p* = 0.139, mean = -0.014, SD = 0.026) did not show such an alignment. One possibility is that having many more onsets and offsets of words than the number of neural state boundaries could explain this difference between words and clauses. We therefore performed a post-hoc analysis in which we extracted the non-optimal GSBS-solution that had the same number of neural state boundaries as the number of onsets and offsets of words, but there was still no significant alignment (low-level ROI: *p* = 0.99; high-level ROI: *p* = 0.98; results not visualized). Together, these results indicate that neural states in ECoG data are indeed aligned with the stimulus, specifically with the on- and offset of clauses. The lack of alignment with words could be due to the electrode locations being too widespread in ECoG to capture such a relatively low-level language feature.

**Fig. 1.**
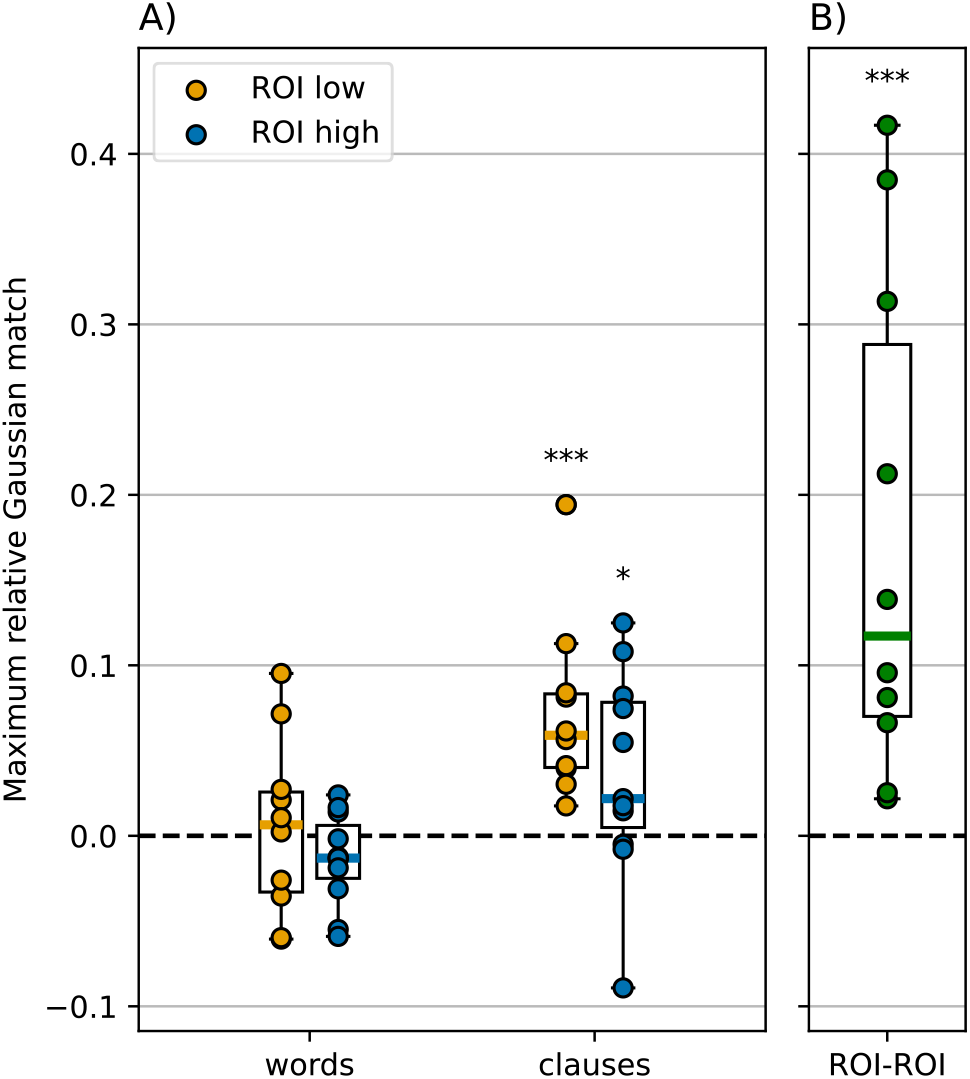
Neural state boundaries in both low- and high-level ROI aligned with clauses but not with words. Additionally, neural state boundaries between the low-level and the high-level ROI significantly align with each other, with each subject having a match above chance. A) Maximum relative Gaussian match per subject for words and clauses, and in both ROIs. Within one box plot, each dot is one subject. B) Maximum relative Gaussian match per subject between the two ROIs. Each dot is one subject. A and B) * *p* < 0.05, ** *p* < 0.01, *** *p* < 0.001.

### Neural boundaries between areas overlap

Based on previous studies with fMRI data, we expected to observe significant overlap between the boundaries in the different ROIs (Baldassano et al., 2017; Geerligs et al., 2022). To investigate this boundary alignment, we computed the maximum relative Gaussian match between the low-level and high-level ROI per subject (Figure 1B). We found that all individual subjects had a maximum relative Gaussian match of above chance-level. Using a one-tailed Wilcoxon signed-rank test on these maximum relative Gaussian matches per subject (N=10), we found that the match between the neural state boundaries of the two ROIs was significantly above zero (*p* < 0.001). This indicates that boundaries indeed showed significant overlap between the two ROIs.

### Duration hierarchy is absent

Previous literature has found a relatively consistent neural state duration hierarchy in fMRI data: low-level areas have shorter neural states than high-level areas (Baldassano et al., 2017), forming a gradient across the cortex (Geerligs et al., 2022). We accordingly expected our high-level language ROI to have longer neural states than the low-level language ROI, and we therefore computed the median neural state length per subject per ROI, taking all speech blocks together. However, we found very inconsistent results across subjects (N = 10): 6 subjects had longer neural states in the low-level ROI, and 4 subjects had longer neural states in the high-level ROI (Figure 2). To investigate the possibility that this inconsistency could be due to taking many anatomically separated Brodmann areas together to define the high-level language ROI, we re-ran GSBS and the analysis on smaller subsets of high-level electrodes that were anatomically close to each other in Supplementary Section C. However, the inconsistency remained and we therefore conclude that the neural state duration hierarchy is not present. This inconsistency with previous literature could perhaps be due to the size and shape of the ROIs. Previous fMRI studies have used small clusters of voxels, while our ECoG electrodes within one ROI encompass a much larger area of 5 or 7 Brodmann areas. Extracting neural states from such a large area could affect the detected state durations in different ways. On the one hand it could diminish neural state boundaries in various smaller areas within the ROI, creating longer neural states overall. On the other hand it could merge multiple neural state boundaries from smaller areas together, creating shorter neural states overall. Alternatively, perhaps the hierarchy of neural state durations is only observable at the timescale of fMRI and not that of electrophysiology. Additionally, previous studies that demonstrated this duration hierarchy of neural states were based on group-averaged data, which makes the state boundaries fully stimulus driven (Baldassano et al., 2017; Geerligs et al., 2022). In contrast, for this dataset, boundaries could have also been driven by endogenous factors, such as mind wandering. This may result in a less robust duration hierarchy.

**Fig. 2.**
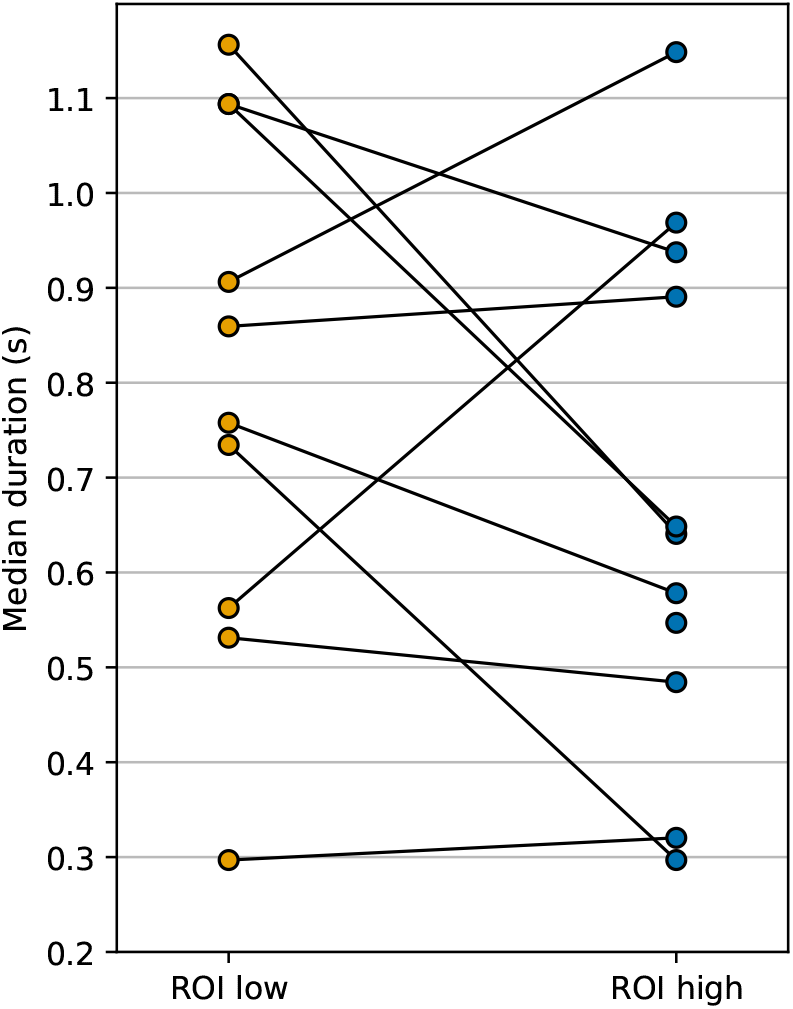
Median state duration in the low and high level ROIs. The durations of neural states are longer in the higher-level ROI than in the low-level ROI for 4 subjects, but longer in the low-level ROI for 6 subjects. Each line is one subject.

### Optimal delay with speech is longer in high-level areas

To investigate the presence of top-down and bottom-up propagation of neural state boundaries, we investigated whether the optimal delay between the stimulus and the neural state boundaries would be different for the two ROIs. The low-level ROI having a shorter optimal delay with speech than the high-level ROI would imply that speech-related boundaries arrive in the low-level ROI before they arrive in the high-level ROI, indicating a bottom-up information flow. Alternatively, the high-level ROI having a shorter optimal delay with speech would be indicative of top-down processes, as speech-related boundaries would then occur in the high-level ROI before they occur in the low-level ROI. Although we had previously shown that the maximum relative Gaussian match between neural state boundaries and clauses was significantly above chance-level at the group level, the subject-level curves of match over delay, as well as the optimal delays, highly varied across subjects (Figure 3A and B). To test whether the optimal delays differed between the low-level and the high-level ROI, we subtracted the optimal delay with clauses in the low-level ROI from the optimal delay in the high-level ROI, and applied a two-tailed Wilcoxon signed-rank test. We only applied this analysis to the optimal delays with the onsets and offsets of clauses, and not of words, as we did not observe a significant alignment between neural state boundaries and word onsets and offsets. We additionally only included subjects with an above-chance relative Gaussian match between neural state boundaries and clauses in both ROIs (N = 7), as the optimal delay is meaningless in subjects with an alignment at or below chance-level. We indeed found that the optimal delay is significantly longer in the low-level ROI relative to the high-level ROI (Figure 3C; *p* = 0.016; mean difference = 123 ms, SD = 0.204 ms). Of the 7 included subjects, 5 had a surprisingly fast optimal delay in the high-level ROI of under 180 ms. Given that all included subjects have a shorter optimal delay with clauses in the high-level ROI than in the low-level ROI, we can conclude that neural state boundaries related to the start and/or end of clauses occur in the high-level ROI before occurring in the low-level ROI, and thus that the alignment with clauses capture a top-down flow of information.

**Fig. 3.**
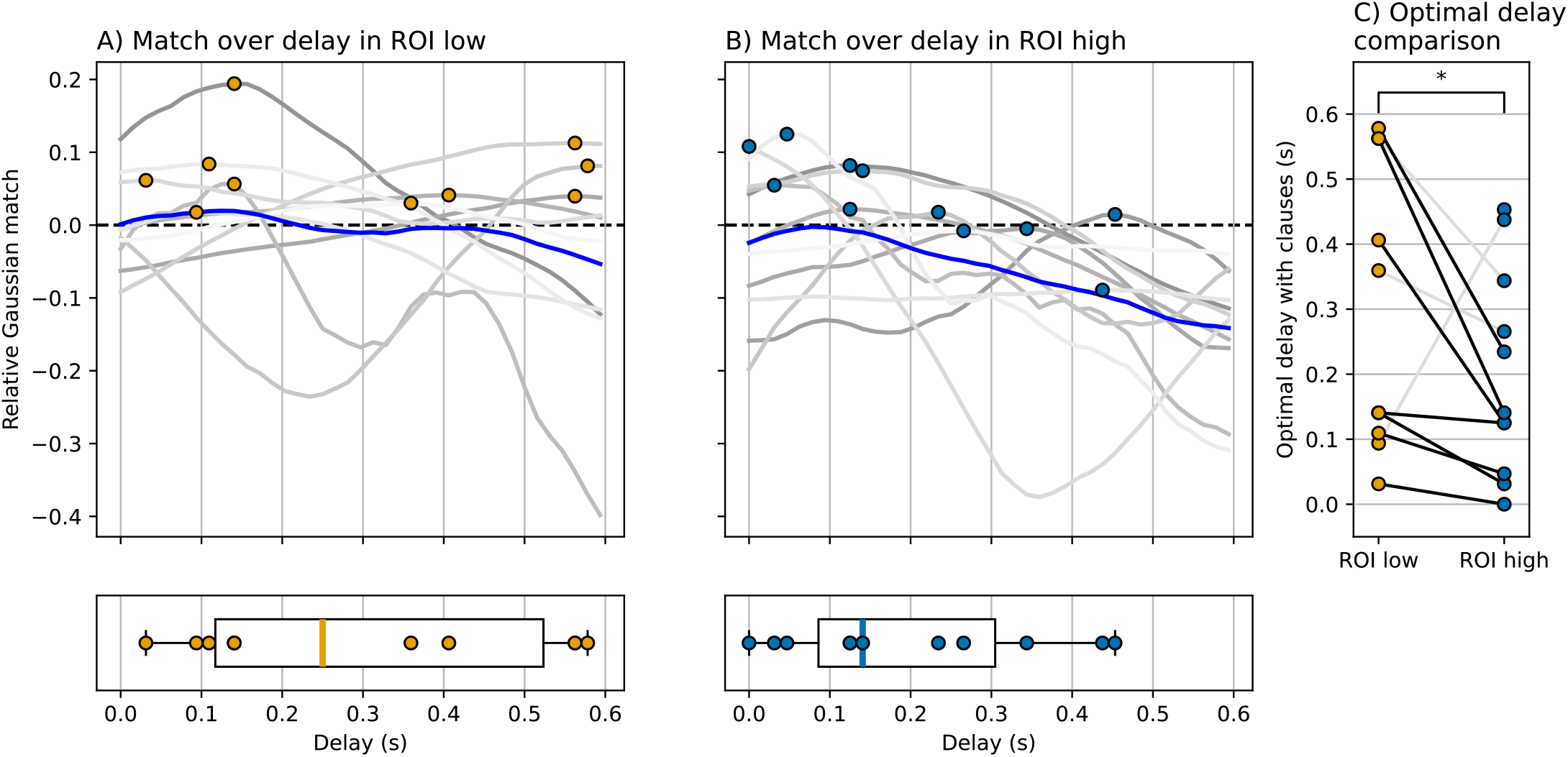
Although the alignment between the onset and offsets of clauses and neural state boundaries was significantly above chance-level (Figure 1A), the optimal delay with those onsets and offsets varied across subjects. When comparing the optimal delays within each subject, the optimal delay with clauses was shorter in the high-level ROI than in the low-level ROI, implying that clause-related neural state boundaries occur in the high-level ROI before the low-level ROI. A) Relative Gaussian match with clauses over delay in the low-level ROI. Each gray line is one subject with a colored marker at the maximum, and the blue line is the average across subjects. B) Relative Gaussian match with clauses over delay in the high-level ROI. C) Optimal delay with clauses per subject. Each line is one subject. Gray lines are subjects with a Gaussian match at or below chance level in at least one ROI. Black lines are subjects with a Gaussian match above chance in both ROIs. * *p* < 0.05.

### Inconsistent delay of boundaries between low- and high-level areas

To investigate whether this top-down flow of information is present across all timepoints or whether it is instead specific to the on- and offsets of clauses, we investigated the delay between boundary timings across the ROIs, irrespective of the stimulus. We did not observe evidence for a consistent delay in the boundary timing across the ROIs (Figure 4). Instead, most subjects had an optimal delay between the two ROIs of around zero with three subjects showing a bottom-up delay (low-level ROI before the high-level ROI), five subjects showing a top-down delay (boundaries in the high-level ROI before the low-level ROI), and two subjects having an optimal delay of exactly 0 s. Given that the alignment with clauses did show a consistent delay, these results suggest that some

**Fig. 4.**
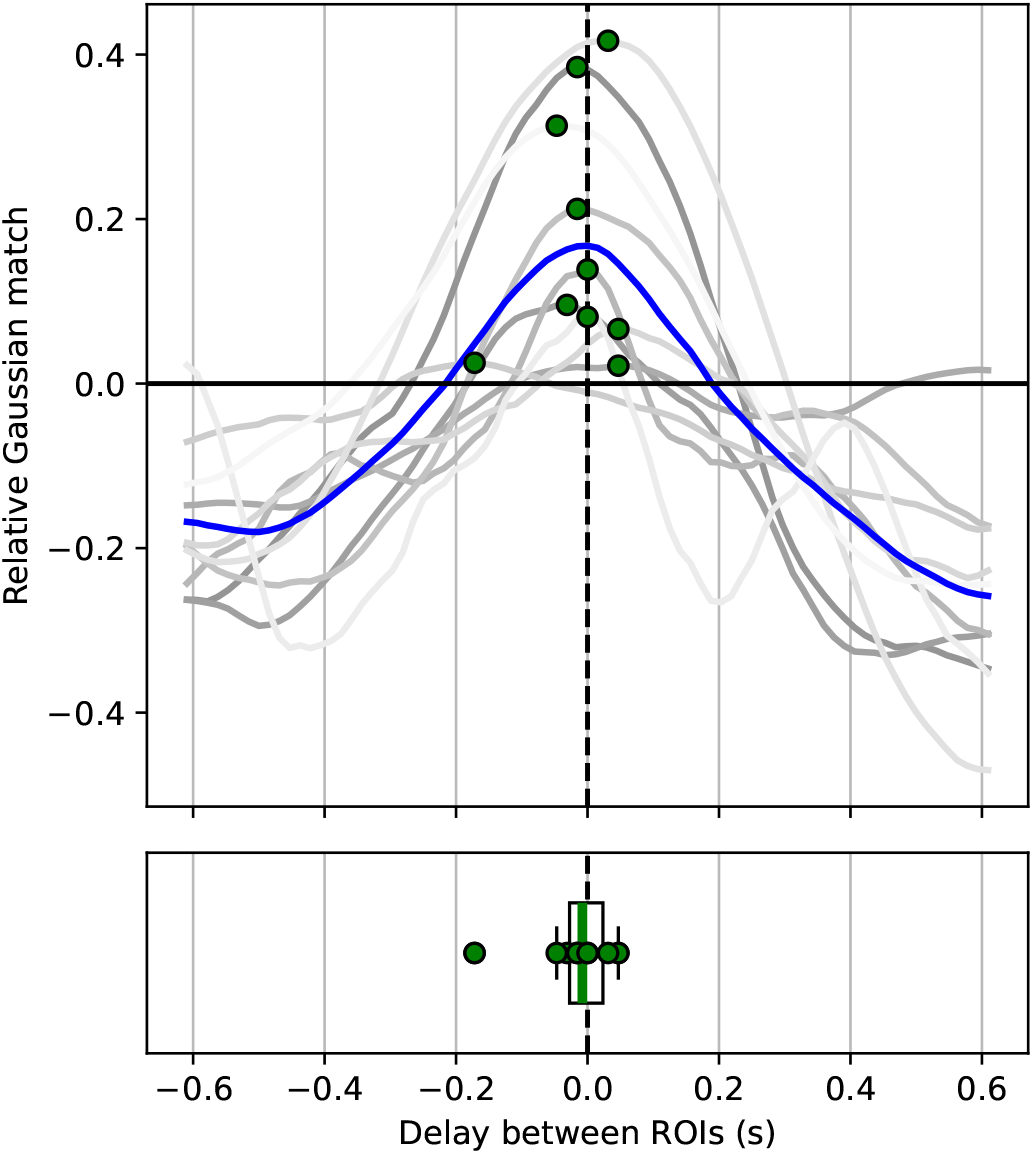
Relative Gaussian match over delay between the low-level and high-level ROI. A negative delay implies that high-level boundaries were generally earlier than low-level boundaries. Each gray line is a subject, and the blue line is the average curve across subjects. The green dots are the maximum relative Gaussian matches per subject.

0.4 boundaries (e.g. those related to higher-level features of the stimulus) may be propagated in a top-down fashion, while other boundaries are propagated in a bottom-up fashion. The latter could include boundaries that are based on more low-level stimulus features, or boundaries that are driven by endogenous changes in brain activity that are not stimulus-driven (e.g., task-irrelevant thoughts or a moment of distraction). Another potential reason for the inter-subject variable delays between the two ROIs could be that different participants had variable electrode locations within each ROI. To sum up, these analyses show that, although the alignment with clauses imply a top-down flow of information, this direction of information flow is not observed across all boundaries.

### Familiarity did not shift neural state boundaries

Having seen the same movie stimulus before affects neural activity patterns. In the context of neural states, Lee et al. (2021) showed that neural state boundaries shift earlier in time the more often subjects had watched the stimulus. As part of the medical procedure, 5 of our subjects had seen the exact same stimulus before, and this could impact our results. To ensure that any difference in boundary timing is not due to multiple exposures, we studied whether the familiarity impacted boundary timing. This analysis is discussed in Supplementary Section D, together with optimal delay results if we were to divide the subjects into two groups based on familiarity. In summary, we did not find the anticipation effect in these ECoG data. This could perhaps be due to the amount of time between viewings, which was several days up to several weeks. With such a long period, many details are likely to be forgotten, making the second viewing less predictable than when viewed on the same day or more than twice. Taking everything together, we conclude that the significant difference in optimal delay with speech between the low-level and high-level ROI (Figure S3B,C,E), as well as the inconsistent optimal delays between ROIs (Figure S3D), cannot be driven by a subset of subjects having watched the stimulus before.

## Discussion

Neural states have previously been investigated as a possible neural underpinning of event cognition. It has been proposed that each area along the cortex segments incoming information into distinct neural states. This proposal is supported by various fMRI and EEG studies that have investigated state boundary alignment with the stimulus as well as the relationship between neural states and memory. Using open ECoG data, here we studied the exact timings of neural state boundaries in multiple brain areas to specifically investigate the presence of bottom-up and topdown propagation of boundaries. We found support for the presence of top-down propagation, extending our understanding of information segmentation throughout the cortex. We additionally replicated multiple attributes of neural states previously observed in fMRI, further supporting the potential of studying neural states through electrophysiological data.

### High temporal resolution uncovers top-down processes

Utilizing the high temporal resolution of ECoG, we studied precise timing differences in neural state boundaries to investigate the presence of top-down and bottom-up processes. Indeed, we found that the optimal delay with the onsets and offsets of clauses is shorter in the high-level than in the low-level area, consistently across subjects. This indicates that clause-relevant neural state boundaries occur in high-level areas before low-level areas.

The exact timing differences we observed between stimulus features and neural states, suggest that the higher-level brain areas may be able to predict the onset and offset of clauses. Previous research has shown that predictability makes neural state boundaries occur earlier in time, and boundaries in higher-level areas are affected more than those in lower-level areas (Lee et al., 2021). Many subjects had an optimal delay of under 180 ms in the high-level language area, while Goldstein et al. (2023) found an optimal delay of 292 ms in the IFG. Additionally, Flinker et al. (2015) reported that the first activation in Broca’s area (part of IFG) in a word-repetition task occurs at 240 ms post stimulus onset, indicating that the clause-related boundaries in our study occur before the IFG has actually received the bottom-up end/start-of-clause information. In the low-level language ROI however, all but one subject had an optimal delay that was longer than that found by previous literature of 55 ms (Goldstein et al., 2023) and 34 ms (Flinker et al., 2015), indicating that the clause-related neural state boundaries occur some time after receiving the relevant bottom-up information. These opposite results in the two ROIs are in line with the finding that, during speech comprehension, temporal regions are involved in bottom-up processing of speech, while frontal regions (including Broca’s area) are more involved in the top-down mechanisms (Zekveld et al., 2006).

Taking these numbers together, we conclude that the onsets and/or offsets of clauses are predicted, leading to neural state boundaries in high-level areas that are earlier in time than what can be expected from a pure bottom-up flow of sensory input. Indeed, language studies have often reported prediction effects during language comprehension, including the possibility of using information from high-level representations to facilitate the processing of incoming information at the level of syntactic structure (e.g., Arai and Keller, 2013) as well as incoming low-level perceptual information (Dikker et al. (2009); see Kuperberg and Jaeger (2016) for a review). Other studies have argued that this prediction could be implemented through a top-down signal that leads to pre-activation (e.g., lexical pre-activation (Fruchter et al., 2015), pre-activation of syntactic gender in spoken language comprehension (Van Berkum et al., 2005), and semantic pre-activation (Altmann and Mirkovic, 2009)). Such pre-activation in high-level language areas, perhaps initiated by higher-level multimodal areas during movie watching, could result in an earlier neural state boundary that relates to the onset or offset of a clause. Although we were only able to show such top-down processes at one level of the cortical hierarchy due to the constraints in this particular study, such processes could be present at other levels as well, creating a hierarchy of linguistic predictions, shown to be present in natural language comprehension using EEG and MEG (Heilbron et al., 2022) and in fMRI (Caucheteux et al., 2023; Schmitt et al., 2021). This prediction hierarchy could go beyond language comprehension and instead encompass information processing in general in a multi-modal naturalistic environment (Schmitt et al., 2021), in line with the idea of event segmentation and a hierarchy of neural states across the cortex (Baldassano et al., 2017).

### Non-continuous top-down processing may indicate global updating of events

Although we did find support for top-down processes at the onsets and offsets of clauses, such processes may not be continuous. Namely, we did not find a consistent delay between the high- and low-level ROI when we looked at all neural state boundaries, irrespective of the stimulus. This suggests that the top-down direction of boundary propagation does not generalize to all boundaries. Together, our results imply that top-down processes happen at or around the onsets and offsets of clauses, rather than all the time. When we generalize this observation beyond our specific stimulus, these results suggest that boundaries can propagate in a top-down manner when there are relevant changes in context that could affect the processing of information in lower-level areas. This is in line with some studies finding bottom-up processes to be dominant (Chang et al., 2022), while others find top-down processes during natural language comprehension (Heilbron et al., 2022).

Given that the top-down processes seem to happen mostly at clauses and not across all neural state boundaries, the flow of information may depend on what exactly is happening in the stimulus. This is an extension of Baldassano et al. (2017)’s proposal in which all brain areas across the cortex segment incoming information, and mostly included upward propagation of information. Our results suggest that a neural state boundary in a high-level area can trigger a top-down flow of information, strengthening information separation in lower-level areas.

In relation to the cognitive skill of event segmentation, previous studies have related event boundaries to group-level neural state boundaries in high-level areas (such as the lateral and medial prefrontal cortex) while participants were watching a movie (Baldassano et al., 2017; Geerligs et al., 2022; Oetringer et al., 2025), or listening to music (Williams et al., 2022), as well as to individual-level neural state boundaries (Sava-Segal et al., 2023). Such event boundaries could then perhaps trigger a top-down updating process in lower-level areas, possibly leading to a change in representation of lower-level features, or a prediction of a change. This would be in line with global event updating: the representation of an event as a whole is updated, even when some features (such as characters or spatial locations) remain stable, akin to Event Segmentation Theory (Zacks et al., 2007). Others have proposed that events could instead be updated incrementally (Huff et al., 2014; Zwaan et al., 1995), which would be more congruent with a bottom-up propagation of information. Given that we found evidence for a top-down flow of information but only at particular points in the stimulus rather than continuously, the global and incremental updating perspectives could be combined, with incremental updating happening in general, and global updating at particular moments that fulfill certain criteria. This would correspond to behavioral findings indeed supporting both global and incremental updating (Bailey and Zacks, 2015; Kurby and Zacks, 2012).

### The concept of neural states generalizes from fMRI to electrophysiology

In the context of event segmentation, neural states have previously mostly been studied in fMRI, and strong support has been found for the hypothesis that all brain areas across the brain segment incoming information. To investigate whether neural states can be studied in ECoG data as well, we first studied whether the neural state boundaries in one brain area were aligned with the neural state boundaries in another. We indeed found within-subject alignments between these areas, with all subjects showing a match above chance-level. This is in line with Baldassano et al. (2017) and Geerligs et al. (2022), who both showed that neural state boundaries are often shared across brain areas.

We additionally demonstrated that our two language areas both showed an alignment with speech in the external environment, and in particular with the onsets and offsets of clauses. This corresponds with the notion that each brain area along the cortex segments information in a way that is at least partially stimulus-driven. This has also been discussed and supported in our previous work (Oetringer et al., 2025), showing that location-sensitive areas tend to have a neural state boundary when there is a change in location in the movie stimulus. We did not find such an alignment with words in the language ROIs, which may be due to the overall size of these regions being too large to capture such low-level characteristics.

One phenomenon that is consistently observed in fMRI, but that we could not replicate with ECoG data, is the hierarchy of neural state durations (Baldassano et al., 2017; Geerligs et al., 2022; Sava-Segal et al., 2023). This may have been because of how we defined our regions of interest, which were relatively large compared to fMRI studies due to the nature of ECoG data. Taking multiple areas together, which each have their own intrinsic timescale, could make the duration hierarchy less robust. We also encountered some challenges with defining the optimal number of neural states, compared to previous fMRI studies. In particular, we found larger intra-subject variability than would be expected, potentially due to the signal-to-noise ratio being lower than in group-averaged fMRI data on which GSBS has been developed and tested (Geerligs et al., 2021). Group-averaging is unfortunately not possible in ECoG due to the subject-specific electrode locations. Although some adjustments were necessary in determining the optimal number of neural states (see Methods), we were still able to extract neural states in a fully data-driven approach. Even though our two ECoG ROIs were larger and much more sparsely sampled than voxels usually studied with fMRI, we were still able to detect neural states, similar to Henderson et al. (2025), Mariola et al. (2022), and Silva et al. (2019), who defined neural states in EEG data. This is not surprising, as there is a large body of literature around neural states at almost all spatiotemporal scales. If we consider neural states to be temporarily stable neural activity patterns that abruptly shift into new patterns regardless of their spatial scale, then neural states have actually been studied with a multitude of imaging methods at a wide range of spatial scales. These include local populations with multivariate single cell activity (Abeles et al., 1995), the whole brain as different configurations of global functional connectivity networks (Song et al., 2023; Yamashita et al., 2021), as well as multivariate whole-brain voxel activity states with fMRI (Zhang et al., 2021), and as microstates with EEG (Michel and Koenig, 2018) and MEG (Tait and Zhang, 2022). In fact, evidence also suggests a direct correspondence between electrophysiological microstates and fMRI functional connectivity states (Britz et al., 2010; Michalopoulos and Bourbakis, 2015). Such “neural states” have been associated with a lot of behavioral and cognitive processes, such as attention, memory, exploration, and more (Artoni et al., 2023; Panagiotides et al., 2011; Song et al., 2023; Suhail et al., 2022; Yamashita et al., 2021). This background makes us confident that our sparsely sampled ECoG neural states are still methodologically sound and behaviorally meaningful.

Although studying neural states in ECoG comes with extra challenges, we were still able to replicate multiple fMRI findings, indicating that neural states are present and relevant in ECoG data as well despite the difference in temporal resolution. In particular, this is the first study to our knowledge that shows a neural state boundary alignment between multiple brain areas in electrophysiological data. Showing this alignment in ECoG is of particular importance, as it cannot be studied in MEG and EEG due to their lower spatial specificity. This study therefore provides a stronger foundation for studying neural states in existing and future electrophysiological studies, including more accessible methods such as EEG and MEG.

## Conclusion

By investigating neural states in ECoG data while subjects were watching a movie, we were able to show that neural state boundaries can be propagated in a top-down manner. In particular, the top-down processes in general language areas take place at onsets and offsets of clauses. We did not find a similar effect for words rather than clauses, possibly because of the electrode placement and subsequent ROI definitions. We additionally showed that fMRI findings generalize to electrophysiology, giving stronger incentive to (further) investigate neural states in electrophysiological data, which includes ECoG, but also EEG and MEG.

## Methods

### Data acquisition

To investigate neural states with high temporal resolution and high spatial specificity, we used an open iEEG dataset (Berezutskaya et al., 2022). In this dataset, 46 epilepsy patients were implanted with electrode grids (ECoG; exposed diameter of 2.3 mm, inter-electrode distance of 10mm) and watched a short audiovisual movie. The data were sampled at a rate of 512Hz or higher. Some subjects (additionally) had depth electrodes and/or a high-density grid implanted, but these types of recordings were not used in the current study. More details about data acquisition can be found in Berezutskaya et al. (2022).

### Participants

Only subjects with ECoG recordings were included. We excluded participants that were younger than 18 (20 subjects) to ensure that the developmental differences in younger brains did not affect the reliability of our results. Additionally, we excluded participants who had electrodes only in the non-language hemisphere (5 subjects), or had major issues in their brain anatomy such as tumors (1 subject), missing a substantial amount of brain tissue (2 subjects), or ventricles of an abnormal size indicative of severe atrophy (2 subjects). After preprocessing, an additional 5 subjects were excluded due to the presence of atypical activity based on visual inspection (see Supplementary Section E for visualizations and discussions). Of the remaining subjects (N = 11, aged 19 to 52 years, 4 male, 7 female), six saw the stimulus once, while another five had seen the stimulus before, with several days or weeks between viewings. Analyses were performed on all 11 subjects together.

### Stimulus

Subjects watched an audiovisual movie of 6.5 minutes that was originally Swedish but dubbed into Dutch. They were free to move their eyes. The movie consisted of 7 music blocks, in which background music was presented and no character was talking, and 6 speech blocks in which the movie was presented as usual, including characters having conversations. These block types were interleaved, starting with music, and each block was 30 seconds. In this study, we only used the speech blocks in our analyses as our analyses focused on language. The dataset additionally included detailed speech annotations of various language features, of which we used clauses and words. A “boundary timeline” for each of these features was created by having 1’s when there was an onset and/or offset of a clause/word, and 0’s at any other timepoint. For words, there were 540 1’s in total, of which 94 (= 17.4%) were offsets without the onset of a new word at the exact same time. For clauses, there were 153 1’s in total, of which 61 (= 39.9%) were offsets of a clause without the onset of a new clause.

### ROI definitions

To be able to investigate the hierarchy of neural states within the language network, we defined one high-level language ROI and one low-level language ROI per subject. Specifically, we divided the language network as defined in Heilbron et al. (2022) into two parts. The high-level ROI consisted of Brodmann areas 38, 39, 40, 44, 45, 46, 47, which include Broca’s area and temporal pole. The low-level ROI consisted of Brodmann areas 20, 21, 22, 41, 42, which include the primary auditory cortex. The Brodmann area per electrode was determined by segmenting the subject-level anatomy into Brodmann areas using the PALS-B12 atlas (Van Essen, 2005) in Freesurfer (version 7.3.2). See Supplementary Section A for the number of electrodes per ROI per subject. One subject did not have any electrodes in the low-level ROI, and was thus excluded from any analysis that required this ROI.

### Preprocessing

To clean the data, we first applied a procedure based on Zada et al. (2024) and Goldstein et al. (2022) in all electrodes within a subject. We first dropped the bad electrodes as labeled by Berezutskaya et al. (2022), who visually inspected the data with respect to artifacts and outliers and checked whether any electrodes were placed on top of others. Then the data was high-pass filtered at 0.1 Hz. Spikes in the data were automatically detected and interpolated. Then, data was re-referenced using either a common average filter or an independent component analysis (ICA) based method, depending on the noise profile of the subject. In case of ICA-based re-referencing, we used the InfoMax algorithm and omitted bad spikes during fitting. Bad components were then selected based on visual inspection: first the mixing matrix was inspected to identify components that were shared among many electrodes, and then the timeseries of these components were inspected and identified as “bad” if it indeed looked non-neural. After re-referencing, a notch filter with a width of 2 Hz was applied at 50 Hz to remove line noise. Then the electrodes were selected per ROI as described above. Additional bad electrodes were identified based on visual inspection of the timecourses of that particular ROI and excluded from analysis. The data was then downsampled to 64 Hz, which was a balance between improving computational complexity and keeping a high enough temporal resolution for our analyses, then clipped at 3 times the standard deviation, and finally z-scored to have a mean of zero for each electrode.

### Data-driven detection of neural state boundaries

To identify the exact locations of neural state boundaries, we used Greedy State Boundary Search (GSBS; Geerligs et al., 2021) on single-subject data, per speech block and per ROI. Given time by space data, GSBS defines neural states iteratively in a fully data-driven manner. In most research so far, GSBS is applied to group-averaged fMRI data, but ECoG data cannot be averaged across subjects due to the subject-specific electrode locations. Notably, previous literature has demonstrated that GSBS works for single-subject electrophysiological data as well (Henderson et al., 2025; Mariola et al., 2022). The parameters were configured according to the original recommendations by Geerligs et al. (2021): the maximum number of neural states was set to half the number of time points in each run, and fine-tuning was set to 1, allowing each neural state boundary to be adjusted by 1 timepoint after each iteration for a better fit if necessary. We specifically utilized the states-GSBS algorithm, which is an enhanced and more reliable version of GSBS introduced by Geerligs et al. (2022), capable of potentially placing a full neural state (i.e., two boundaries) in a single iteration instead of always placing just one boundary at a time.

Within GSBS, the optimal number of states is determined using the t-distance metric, which represents how similar within-state activity patterns are and how dissimilar activity patterns between consecutive states are. The optimal number of states is then determined by taking the number of states that gives the highest t-distance. However, we have found that this approach results in the number of states being highly variable across blocks in this particular dataset, with for example one subject having 20 to 31 states per 30 seconds for 5 blocks, and having 152 states for the remaining 30-second block. This discrepancy may be because of the lower signal-to-noise ratio in single-subject ECoG data, as compared to group-averaged fMRI data for which GSBS was originally developed and tested. To achieve a more stable state segmentation across blocks within each participant, we adjusted our procedure for selecting the optimal number of states. First, we averaged the t-distance metric across blocks, resulting in an average optimal number of states per 30 seconds by taking the highest t-distance from this averaged distribution. Next, we fine-tuned the number of states per block, by finding a peak in the t-distance metric per block that is closest to the chosen optimal number of states across all blocks. Here, a peak is defined as any point, or a pair of points with the exact same value, with a lower value on both the left and the right side.

### Gaussian match

To determine the extent of having a “match” between two boundary timelines, such as when comparing neural state boundaries in one ROI to the other, we developed a new metric called Gaussian match. Previously used match metrics applied to fMRI data such as counting the number of overlapping boundaries (Baldassano et al., 2017; Geerligs et al., 2022) or correlations (Oetringer et al., 2025) are not applicable here, since the much higher temporal resolution of ECoG data requires more flexibility in the relative timing of two matching boundaries. We therefore developed a measure that weighs the amount of match depending on the temporal distance between two boundaries, rather than having an all-or-nothing approach. We did this by projecting a Gaussian (SD = 332 ms, mean of 0), of which the amplitude has been scaled between 0 and 1, onto each boundary in one timeline. Then, to compute the match of the current boundary, the closest boundary in the other timeline was selected and the match was determined using the amplitude of the projected Gaussian at the timepoint of this closest boundary. Thus, the further away the closest boundary in the other timeline is located, the lower the match, with a maximum of 1 when the two boundaries occur at the exact same point in time.

To determine the overall match between two timelines, one timeline is chosen as the seed. Then for each boundary in this seed timeline, the match is computed using the closest boundary in the other timeline. Then, the average match across all boundaries is computed as the Gaussian match. Thus, the Gaussian match metric is non-commutative, since choosing the other timeline as the seed gives a different Gaussian match value. Therefore, it is important to consider which timeline becomes the seed timeline. When investigating whether boundaries in stimulus features co-occur with neural state boundaries, we used the stimulus feature timeline as seed since having extra neural state boundaries should not have a negative impact on the match. When comparing neural state boundaries between different ROIs, we took the timeline with the lowest number of boundaries as the seed boundary, to prevent one area having more boundaries than the other from negatively impacting the match value.

### Accounting for delays

Because of the high temporal resolution of electrophysiological data, a delay may be detected between the stimulus and corresponding activity in a specific brain area, or between two brain areas. Such delays have been estimated before in ECoG (Flinker et al., 2015), with an onset as early as 34 ms in the temporal cortex and 240 ms in Broca’s area, but the exact delay will depend on the specific electrode locations which differ per subject. When computing the match between two timelines, we therefore computed the Gaussian match over various delays. When comparing stimulus features to neural features, we shifted the stimulus feature timeline forward in time ranging from a delay of 0 ms to 600 ms, with steps of 1/64 s (*≈* 16 ms). When comparing two timelines of neural features, the possible delays instead ranged from roughly -600 ms to 600 ms, with steps of 1/64 s. The optimal delay was then defined as the delay that gave the maximum Gaussian match.

### Relative Gaussian match

The Gaussian match in the data (*match*_*data*) will always be above zero, also when boundaries are not aligned with each other. The Gaussian match that can be expected by chance is dependent on the number of neural states and their durations in each timeline. In order to get a measure for the match relative to chance-level, we computed a null distribution of the maximum Gaussian match by shuffling the neural states in one timeline 1000 times, and recomputing the maximum Gaussian match over all delays per permutation. The chance-level Gaussian match is then defined by taking the mean of all the maximum Gaussian matches across permutations (giving us *match*_*null*). The relative Gaussian match was then defined as (*match*_*data −match*_*null*)*/*(1*− match*_*null*), giving a value of around 0 when the Gaussian match in the data is at chance-level, and 1 when this match is perfect. Group-level statistics can then be computed over the maximum relative Gaussian match per subject using the Wilcoxon signed-rank test.

## Data and code availability

The ECoG data and stimulus annotations were published by Berezutskaya et al. (2022) and are freely available at openneuro.org/datasets/ds003688. All Python code for further data preprocessing and analysis can be found at github.com/dynac-lab/temporal_propagation.

## Ethics

This study utilized open ECoG data. According to the accompanying literature (Berezutskaya et al., 2022), all participants were admitted to the University Medical Center Utrecht, Netherlands, for diagnostic procedures related to epilepsy. For some participants, the movie stimulus with ECoG measurements were part of clinical function mapping procedures, and these participants gave a written permission to use their clinical data for research purposes. Others participated as part of scientific research, in which case they gave their written informed consent to participate in research tasks. All participants gave their consent to share their de-identified data publicly.

## Contributions

**Djamari Oetringer**: Conceptualization, Methodology, Software, Formal analysis, Visualization, Writing - Original Draft. **Sarah Henderson**: Conceptualization, Methodology, Writing - Review & Editing. **Dora Gözükara**: Methodology, Writing - Review & Editing. **Linda Geerligs**: Conceptualization, Methodology, Validation, Writing - Review & Editing, Funding acquisition, Supervision.

## Funding

Linda Geerligs was supported by a Vidi grant (VI.Vidi.201.150) from the Netherlands Organization for Scientific Research.

## Declaration of Competing Interests

The authors declare that no competing interests exist.

## Acknowledgments

We would like to thank Julia Berezutskaya and Emily Mo Nipshagen for fruitful discussions and tips on investigating ECoG data, and Mesian Tilmatine and Sahel Azizpour for helpful insights on language processing. We additionally want to thank Umut Güçlü for discussions and insights during the development of this study and for providing feedback on our manuscript.

## Supplementary materials

### A Subject descriptions

The number of electrodes used in the analyses per subject per ROI can be found in the table below. Note that subject number 12 had no electrodes in the low-level ROI and was therefore excluded from any of the analyses that required this ROI. 6 subjects viewed the stimulus only once (“Novel”), while 5 subjects (“Familiar”) had seen the stimulus before.

**Table S1.**
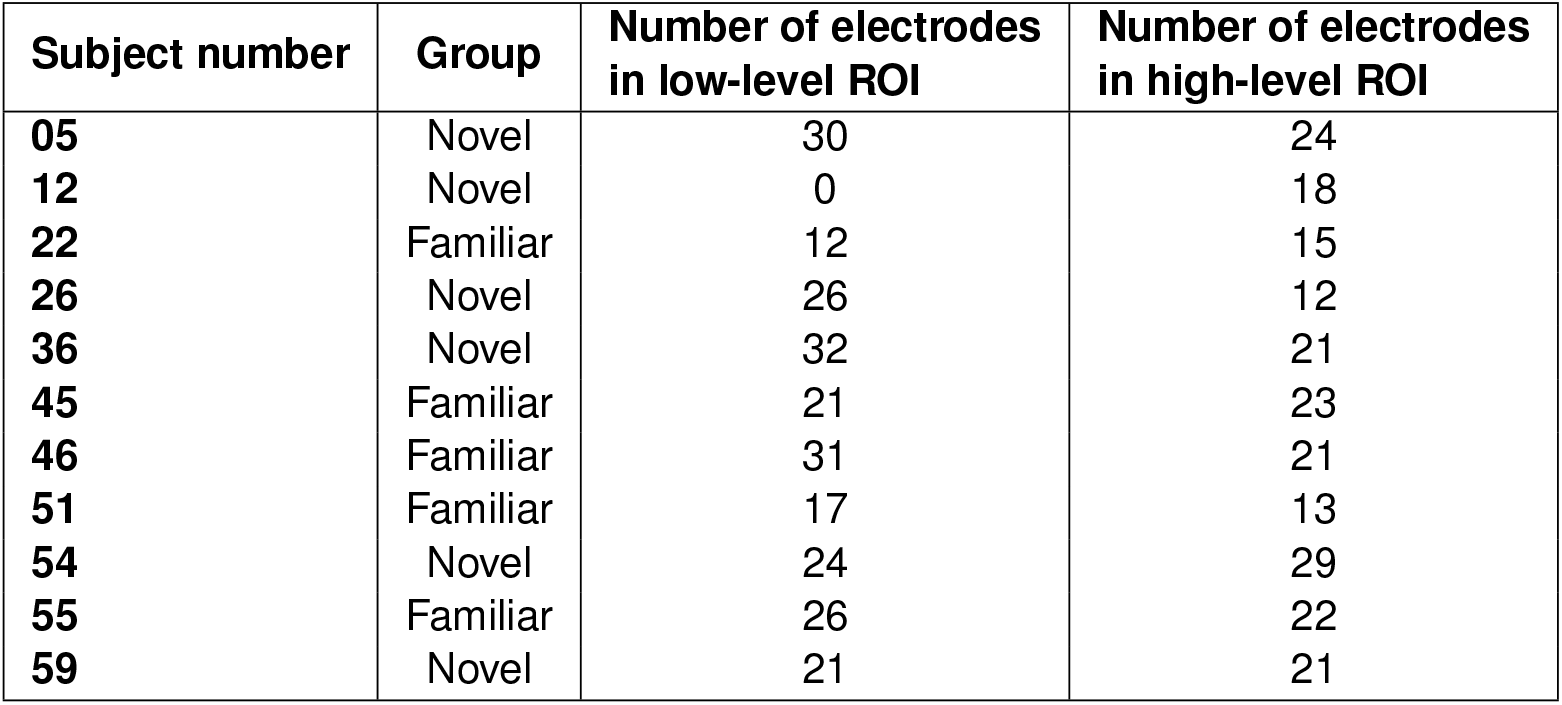
Summary descriptions of the used electrodes per subject.

| Subject number | Group | Number of electrodes in low-level ROI | Number of electrodes in high-level ROI |
| --- | --- | --- | --- |
| 05 | Novel | 30 | 24 |
| 12 | Novel | 0 | 18 |
| 22 | Familiar | 12 | 15 |
| 26 | Novel | 26 | 12 |
| 36 | Novel | 32 | 21 |
| 45 | Familiar | 21 | 23 |
| 46 | Familiar | 31 | 21 |
| 51 | Familiar | 17 | 13 |
| 54 | Novel | 24 | 29 |
| 55 | Familiar | 26 | 22 |
| 59 | Novel | 21 | 21 |

### B Example timelines of neural state boundaries

**Fig. S1.**
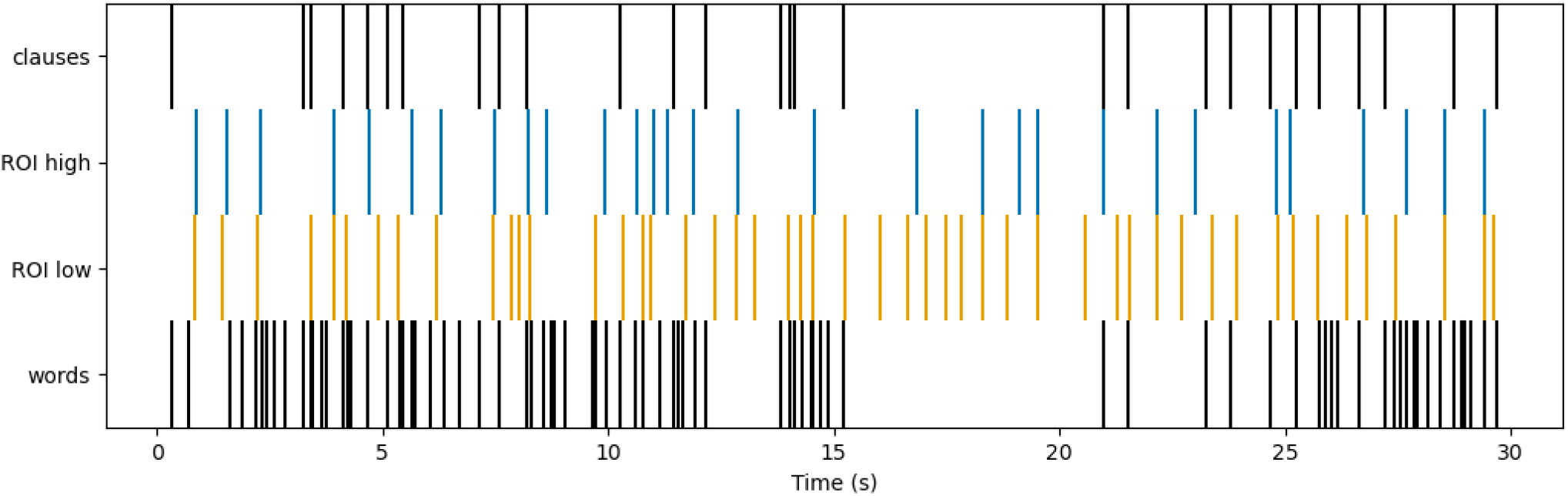
Timeline of the onsets and offsets of clauses and words of one example block, together with the neural state boundaries in the low- and high-level ROI of one example subject. No delay has been applied.

The timings of annotation boundaries of an example run are visualized in Figure S1, together with the neural state boundaries in both the low-level and high-level ROI of one example subject.

### C Duration hierarchy using smaller high-level ROIs

When investigating the neural state durations in our main analysis, we expected the high-level ROI to have longer states than the low-level ROI. However, the results were inconsistent across subjects with 6 subjects having longer neural states in the low-level ROI, and 4 subjects having longer neural states in the high-level ROI. Given that the high-level ROI consists of multiple anatomically separated Brodmann areas, here we studied whether applying GSBS and subsequent analysis on smaller and anatomically connected areas would give rise to more consistent results. Therefore, we divided the Brodmann areas up into 3 smaller areas: temporal pole (TP; Brodmann area 38), angular gyrus and connected areas (AG+; Brodmann areas 39 and 40), and Broca and connected areas (Broca+; Brodmann areas 44, 45, 46, and 47). We re-ran GSBS and subsequent analyses per subject only if the number of electrodes in a smaller area was at least 10. If the duration hierarchy was indeed inconsistent in the main analysis because of these anatomically separated areas being taken together, then we expect all of these smaller areas to show longer median durations as compared to the low-level ROI in the same subject. This was however not the case for any of the three areas (Figure S2). TP had the exact same median state duration as the low-level ROI in 1 subject, and shorter states in TP than the low-level ROI in another subject. AG+ had longer states than the low-level ROI in 2 subjects, and shorter states in another 2 subjects. Finally, Broca+ had longer states than the low-level ROI in 2 subjects, and shorter states in another 2 subjects. Taking everything together, we conclude that extracting neural states from anatomically connected Brodmann areas only still does not give rise to the neural state duration hierarchy found in previous fMRI studies. Together with the results of our main analysis, we conclude that the neural state duration hierarchy is absent in these data.

**Fig. S2.**
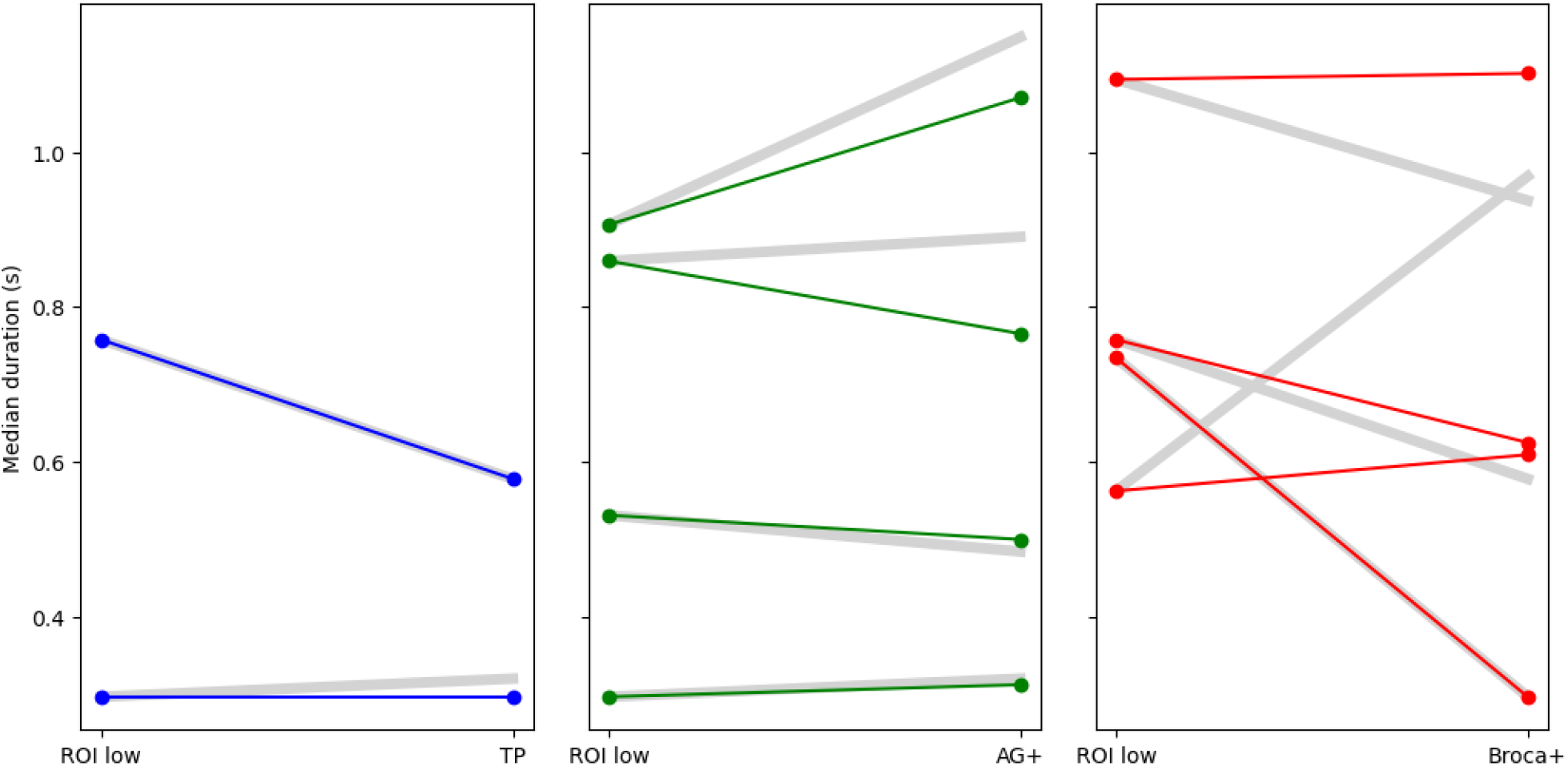
Median state duration in the low-level ROI compared to the high-level ROI, but divided up into three smaller high-level areas: temporal pole (TP), angular gyrus and connected areas (AG+), and Broca and connected areas (Broca+). Each colored line is one subject, with the gray thick line indicating the median durations in the low-level ROI compared to the high-level ROI that was used in the main analysis.

### D Anticipation analysis

To ensure that having seen the stimulus before did not impact the results, we investigated the timing differences of neural state boundaries of subjects who had already seen the stimulus (the Familiar group) to those of the subjects who viewed the stimulus for the first time (the Novel group). More specifically, we tested whether boundaries of the Familiar group occurred earlier in time than those of the Novel group. Per subject pair, we first selected a group of electrodes that the subjects had “in common”. Here, “in common” means that the electrodes were located in the same area according to the Brainnetome Atlas (Fan et al., 2016). If one subject had more electrodes in a particular area than in another, random electrodes of that subject were dropped until the number of electrodes within a particular area was equal between the two subjects. We then ran GSBS on both subjects separately, using all selected electrodes across the brain as input, and computed the optimal delay with clauses as was done in our main analysis. GSBS and consequently the optimal delay was only computed if the number of electrodes that the two subjects had in common was at least 15. By doing this for all subject pairs, we created a subject by subject matrix indicating the difference in optimal delay with clauses for each subject pair (Figure S3A). The measurement to be statistically tested was the median of the lower-left quadrant, which indicated how much later the optimal delay of the familiar subjects was compared to the novel subjects. Based on (Lee et al., 2021), this number is expected to be negative as familiarity moves neural state boundaries earlier in time, making the optimal delay of familiar subjects shorter. Indeed, the median of the lower-left quadrant is equal to -0.008 s. To statistically test whether this number is significantly below 0, we permuted the subject by subject matrix. This was done by shuffling the order of the subjects 10.000 times to create a null distribution, while labeling the first 6 subjects as “Novel” and the last 5 subjects as “Familiar”, as was the case for the original non-permuted data. In the final analysis, the boundaries of Familiar subjects were not earlier in time than those of Novel subjects (p = 0.6331). We therefore conclude that the Novel/Familiar distinction between subjects did not affect boundary timing, and thus that the conclusions of our main analyses still hold.

If an anticipation effect had been present, it could have affected the results of our main analysis regarding optimal delays with speech and between ROIs (Figure 3 and Figure 4 in the main text). These results are presented again in Figure S3B, C, D and E, but now with Novel and Familiar subjects separated. In line with the absence of an anticipation effect, the optimal delays are not substantially different between Novel and Familiar subjects.

**Fig. S3.**
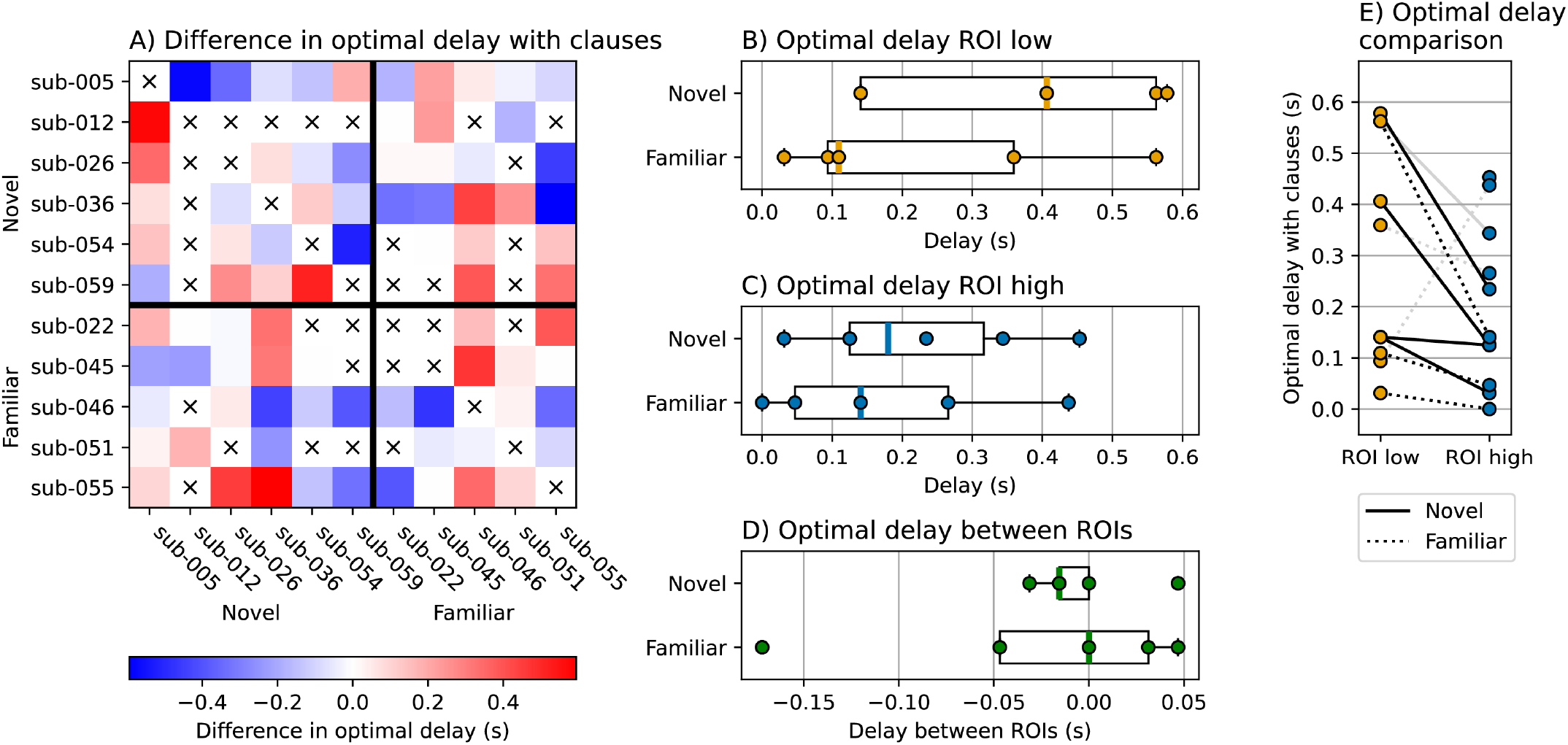
Stimulus familiarity did not affect timing results. A) Difference in optimal delay with clauses between subjects in the Novel and Familiar groups. A negative number (blue) means that the optimal delay with clauses was shorter for the subject on the y-axis than for the subject on the x-axis. A cross indicates that the subjects could not be compared because of having too few electrodes in common. B and C) Optimal delay with clauses in the low-level and the high-level ROIs. These are the same data as in Figure 3A and B, now divided up into the Novel and the Familiar groups. D) Optimal delay between ROIs. These are the same data as in Figure 4, but divided up into the Novel and the Familiar groups. E) Optimal delays with clauses per subject. Each line is one subject. Gray lines are subjects with a Gaussian match at or below chance level in at least one ROI. These are the same data as in Figure 3C, but divided up into the Novel and the Familiar groups.

### E Artifacts in data

After excluding subjects based on age, language hemisphere, and brain anatomy, we visually inspected the timeseries of all electrodes and the time by time correlations of all remaining subjects. As all subjects were patients with severe epilepsy, more atypical brain activity can be expected as compared to healthy controls. Based on these visualizations, another 5 subjects were excluded due to the presence of atypical signals, which we classified as artifacts from unknown sources. For comparison, the data of two included subjects are visualized in Figure S4.

**Fig. S4.**
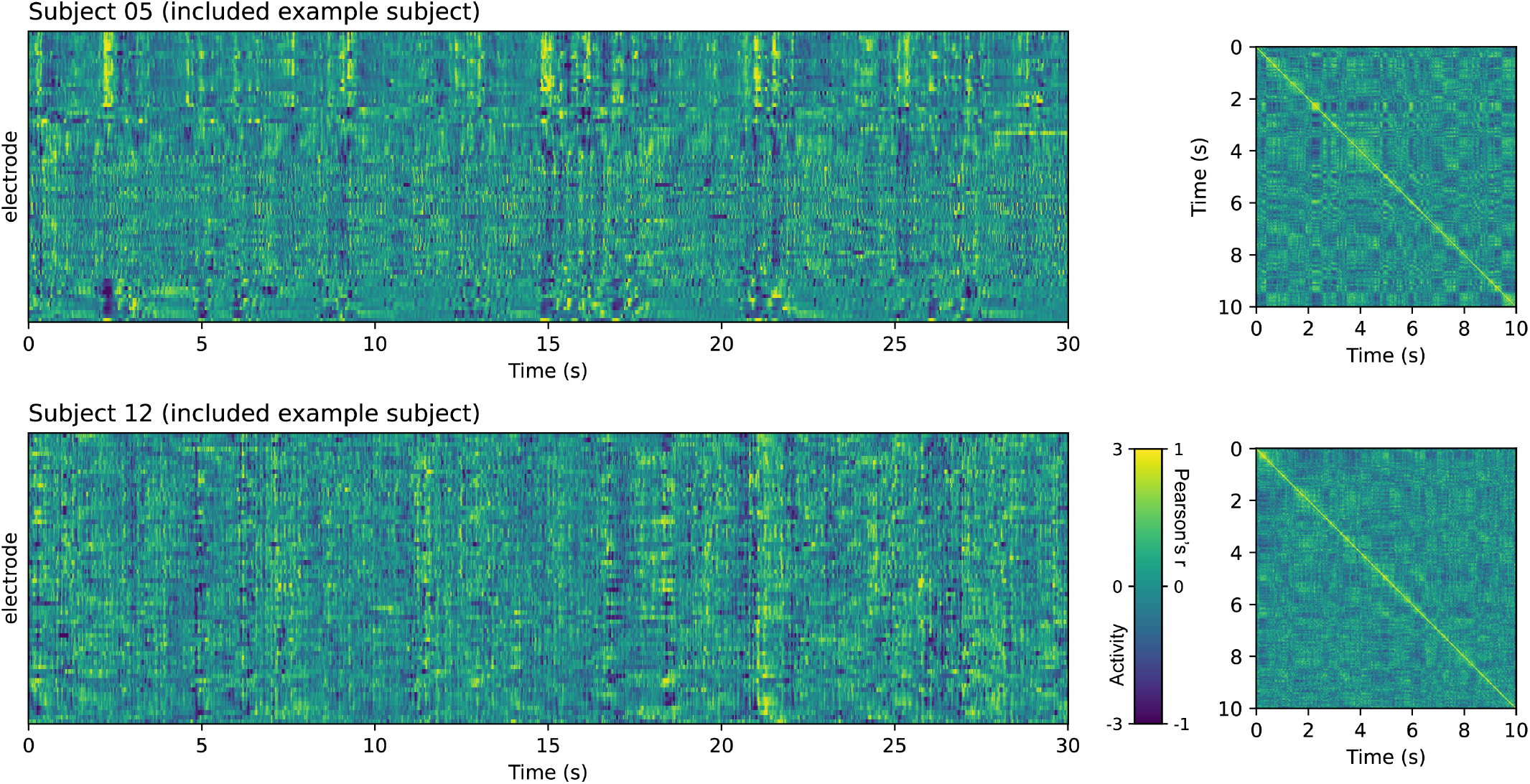
Preprocessed data of one example block for two subjects that were included in the final analyses. Left: timeseries of a full 30-second block. Right: time by time correlation matrices of the first 10 seconds of that block.

For each subject that was excluded on the basis of visual inspection, the preprocessed data of one block of 30 seconds in visualized in Figure S5. Note that the data in this figure was re-referenced using the ICA-based method, but for every depicted subject the artifacts were also present when using the common average rereference.

#### Subject 03 and 07

Subjects 03 and 07 had a rhythmic artifact throughout the whole stimulus, although the rhythm of subject 07 seems to have a higher frequency. The presence of such a rhythm is especially apparent in their time by time correlations, which form a pattern similar to a checkerboard. The durations of the periods with relatively high correlations are more consistent than what can be expected from typical neural activity.

#### Subject 48 and 57

Both subject 48 and 57 had bursts of atypical signal with periods of very high or very low neural activity. This was especially apparent in their timeseries but also reflected in their time by time correlations. This can be seen once in each example block depicted in the figure, but also occurred at other moments throughout the whole stimulus.

#### Subject 60

Many electrodes of subject 60 had non-synchronized atypical activity, which can be seen as wide horizontal stripes of prolonged negative or positive activity in the timeseries. The set of electrodes with such atypical activity differed per block, though there was some overlap. Given the high number of bad electrodes, especially when taking all blocks together, we decided to exclude this subject.

**Fig. S5.**
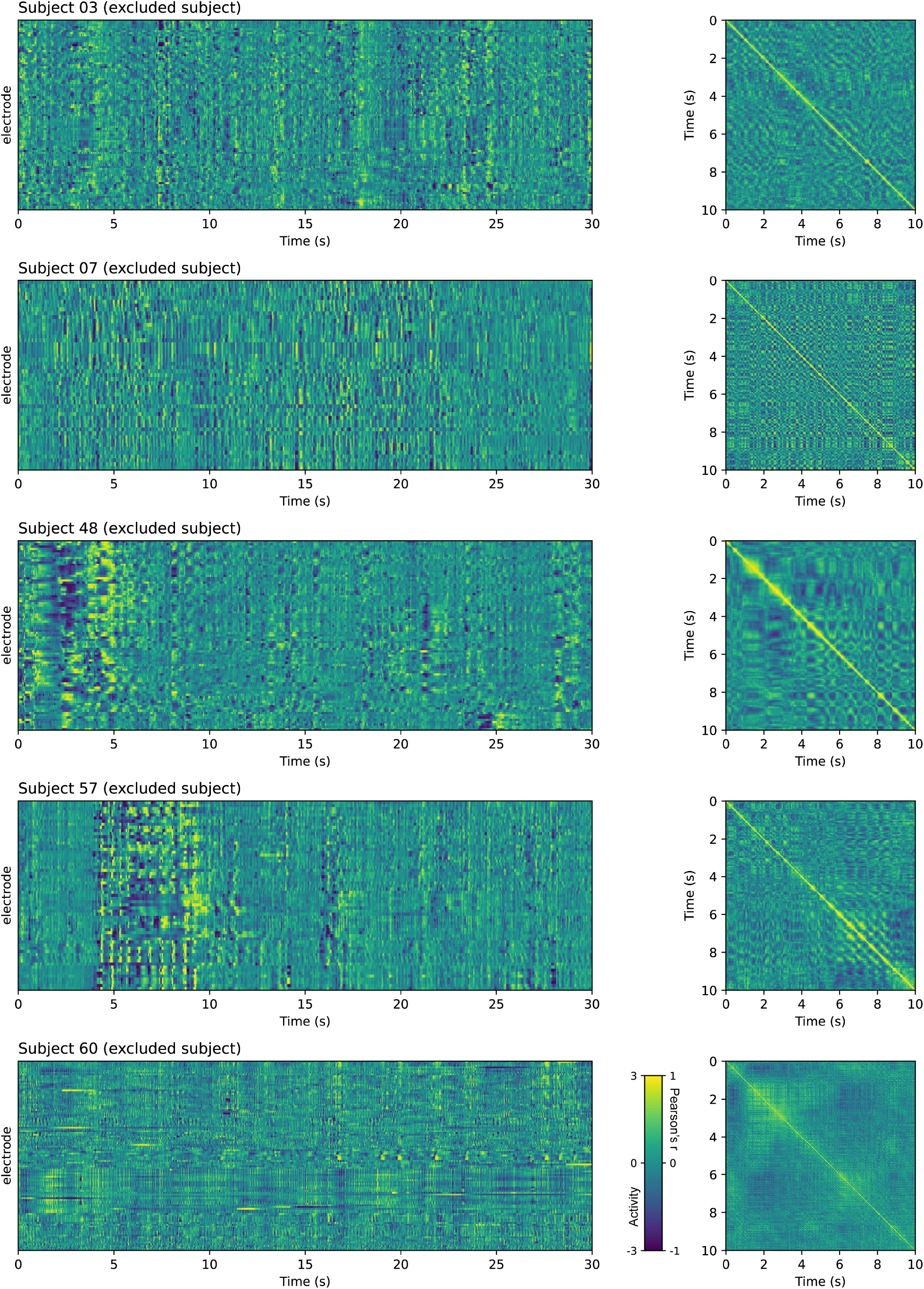
Preprocessed data of one example block for each subject that was excluded from the analyses based on visual inspection of their data. Left: timeseries of the full 30-second block. Right: time by time correlation matrices of the first 10 seconds of that block.

## Notes

https://github.com/dynac-lab/temporal_propagation

### Competing Interest Statement

The authors have declared no competing interest.

### Summary of Updates

A small but crucial typo has been resolved. Namely, the previous version said that we used a Gaussian with a standard deviation of 100ms, while it was actually 332ms. Additionally, some design changes have been applied that do not affect the contents of the manuscript.

